# Fast and sensitive diffuse correlation spectroscopy with highly parallelized single photon detection

**DOI:** 10.1101/2020.07.08.193433

**Authors:** Wenhui Liu, Ruobing Qian, Shiqi Xu, Pavan Chandra Konda, Mark Harfouche, Dawid Borycki, Joakim Jönsson, Edouard Berrocal, Colin Cooke, Haoqian Wang, Qionghai Dai, Roarke W. Horstmeyer

## Abstract

Diffuse correlation spectroscopy (DCS) is a well-established method that measures rapid changes in scattered coherent light to identify blood flow and functional dynamics within tissue. While its sensitivity to minute scatterer displacements leads to a number of unique advantages, conventional DCS systems become photon-limited when attempting to probe deep into tissue, which leads to long measurement windows (∼1 sec). Here, we present a high-sensitivity DCS system with 1024 parallel detection channels integrated within a single-photon avalanche diode (SPAD) array, and demonstrate the ability to detect mm-scale perturbations up to 1 cm deep within a tissue-like phantom at up to 33 Hz sampling rate. We also show that this highly parallelized strategy can measure the human pulse at high fidelity and detect behaviorally-induced physiological variations from above the human prefrontal cortex. By greatly improving detection sensitivity and speed, highly parallelized DCS opens up new experiments for high-speed biological signal measurement.

---

Various optical technologies have been developed to noninvasively detect dynamic biological events in deep tissue, such as changes in blood flow and other related fluctuations. For instance, methods such as diffuse optical tomography (DOT)^1^, diffuse optical spectroscopy (DOS)^2^ and functional near-infrared spectroscopy (fNIRs)^3^ all illuminate a tissue’s surface and measure the resulting scattered light at different locations to reconstruct images of absorption and scattering properties. Recent extensions now utilize time-gated measurements to probe even deeper by isolating scattering from non-superficial layers^4^. By examining scattered light’s spectral interference fringe pattern^5^, the related technique of interferometric near-infrared spectroscopy (iNIRS) can also extract additional path-length-dependent information to accurately measure deep blood flow^6^. The above techniques are distinct from other measurement strategies such as Doppler OCT^7^, OCT angiography^8^ and laser speckle contrast imaging^9^, in that their primary aim is to penetrate multiple millimeters into tissue to extract useful signal.

Another well-established method to measure such deep tissue dynamics, termed diffuse correlation spectroscopy (DCS), utilizes the relatively simple strategy of illuminating tissue with highly coherent light and collecting back-scattered interference using a photon-counting detector. Dynamic events, such as blood flow or other small movements, are then estimated by calculating and processing the scattered light’s temporal autocorrelation curve^10,11^. Unlike the above methods, DCS offers a non-invasive phase-sensitive measurement strategy^12,13^ or deep-tissue motion without requiring a reference beam, and thus offers a promising means to achieve a simple and low-cost functional imaging technique with high sensitivity. DCS has been actively studied and translated into a useful clinical technique for analysis of the brain, breast, and skin^14-18^.

DCS systems are typically implemented in a reflection geometry, where a source and a detector are placed a finite distance *d* apart and detected light is assumed to travel through a “banana-shaped” point-spread function (PSF) that describes the light’s most probable path through scattering tissue^10^ (see Fig. 1). As this PSF’s penetration depth scales linearly with *d*^19^, large source-detector separations are typically required to probe deep within a tissue. However, due to the forward-scattering nature of a biological tissue, most photons do not reach a detector in standard DCS measurement geometries, especially when the source and detector are place a large distance apart to probe deep events. For example, when illuminating tissue with 200 mW/cm^2^ visible light (at the ANSI safety limit^20^), a straightforward simulation^21^ specifies that a single optical mode (i.e., a speckle grain) at a 2 cm source-detector separation includes approximately 10^6^ photons per second (see Supplementary Table. 1). Accordingly, it is possible to detect just one photon, on average, when sampling at a 1 MHz detection rate. Probing variations that may arise beneath the human skull requires large, multi-cm source-detector separations^10,11,22,23^, indicating that current DCS systems are severely photon-limited when outfitted to measure brain dynamics. To partially overcome this low photon budget, current DCS systems must integrate measured signals over a long time period – often more than a second - reducing overall system temporal resolution ^13,24,25.^ While DCS can, in principle, capture the effects of very small non-absorptive changes within the brain, such as cerebral blood flow (CBF)^26^, single-detector implementations of DCS quickly run into an issue of not having enough photons when aiming to detect deep tissue events.

**Fig. 1.**
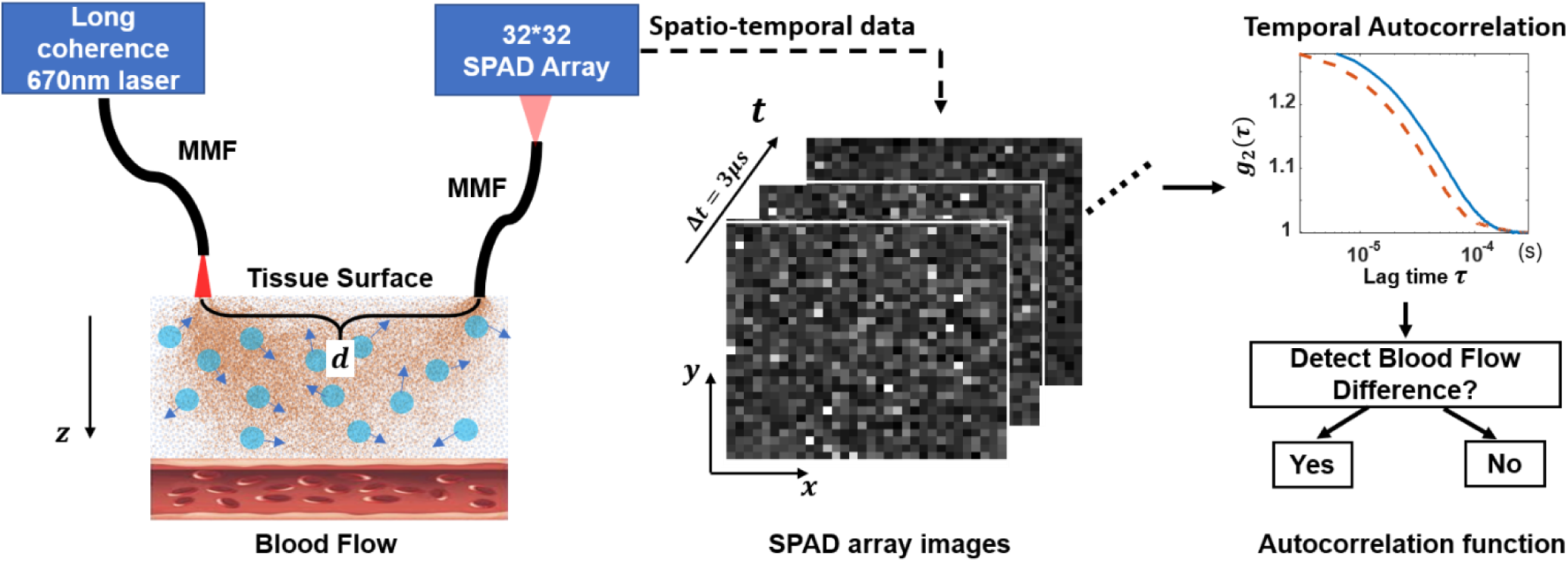
Schematic diagram of parallelized DCS detection. The light from a 670 nm long-coherence laser is delivered by a multimode fiber to the tissue surface. Photons that penetrated deep enough will be reflected by the moving red blood cells and other scatterers. A small fraction will scatter back to the detection multimode fiber and are directed to the SPAD array. The distance between the center of the illumination fiber tip and detection fiber tip is *d*. The SPAD array records the temporal intensity fluctuations of the speckles at a sampling rate of 333 kHz across all 1024 channels. The autocorrelation curves *g*_2_ of each SPAD are computed and combined. Perturbations induced by blood flow are detected by comparing the decorrelation rate of the final averaged result.

A clear solution to this problem is to simply detect light from more optical modes - measuring light from *N* speckle grains yields *N* times more photons, and thus a proportionally improved signal-to-noise ratio (SNR). Since the optical modes emerging from the tissue are nearly mutually incoherent (i.e. decorrelating independently), sampling multiple speckle grains on a single detector reduces measurement contrast and is not a helpful strategy (beyond integrating over a few modes per channel^27,28^). DCS thus inherently benefits from a parallelized detection strategy, wherein multiple independent detectors could be used to measure *N* fluctuating speckle grains simultaneously to extract proportionally more useful signal from deep tissue. Several multi-channel DCS systems have previously been developed to simultaneously detect more signal. For example, a number of 4-channel DCS systems have been reported in prior work to detect blood flow both from humans^29,30^ and mice^31^; Staples et al. constructed an 8-channel DCS system using one PMT and seven single photon counting modules^32^; Johansson et al. developed a 5×5 SPAD configuration to demonstrate improved SNR on a milk phantom and an *in vivo* blood occlusion test^27^, and Dietsche et al. implemented a 28-channel DCS setup with multiple single-mode fibers to measure blood flow in deep tissue^28^. The DCS system in the last work has the most parallelized detection channels that we are aware of to date.

The recent development of integrated SPAD array technology, wherein thousands or more individual SPAD detectors can be integrated on a single CMOS chip, opens up a new regime for massively parallel single-photon detection, without dramatically increasing the complexity of the DCS measurement setup. SPAD arrays are solid-state detectors now fabricated in standard CMOS technology that, when manufactured at scale, offer a low-cost photon counting solution with unrivaled performance via chip or pixel-level processing capabilities^33^. SPAD arrays have recently been implemented in many biological imaging experiments to achieve a wide variety of interesting tasks. These include depth ranging and imaging^34^, fluorescent lifetime imaging microscopy^35-38^, endomicroscopy^39^, spectroscopy^40^, Raman spectroscopy^41-43^, and localization-based super-resolution imaging^44^, to name a few.

In this work, we use an integrated SPAD array that contains 1024 independent detectors^45^ to demonstrate a massively parallelized DCS system with approximately 40 times more channels than any prior work (to the best of our knowledge). We discuss a unique software pipeline to integrate these parallelized measurements, and quantitatively demonstrate the sensitivity and temporal resolution enhancements of parallelized detection with a novel phantom that consists of a digital micromirror device (DMD) placed directly behind a rapidly decorrelating tissue-mimicking phantom (Fig. 2(b)). The DMD is employed to generate high-speed temporal perturbations with known parameters which allows us to carefully measure and assess system performance. This phantom allowed us to determine our parallelized DCS system’s sensitivity (i.e., its smallest detectable perturbation size) through thick tissue phantoms under various conditions (e.g., average scattering mean-free path, source-to-detector separation, perturbation speed, and signal integration time). For example, we demonstrate the ability to detect a 2.7 × 2.7 mm perturbation under 1 cm of tissue-like phantom (*μ*_*a*_ = 0.4 cm^-1^, 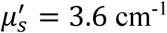) at an autocorrelation curve rate of 0.1 s with 80% accuracy. We also verify that that our system can capture high fidelity *in vivo* blood flow signal from the adult forehead at a 30 ms temporal resolution, and successfully detected an increase in decorrelation time from above the human prefrontal cortex at a similar rate when participants switched between the behavioral tasks of resting versus reading text.

**Fig. 2.**
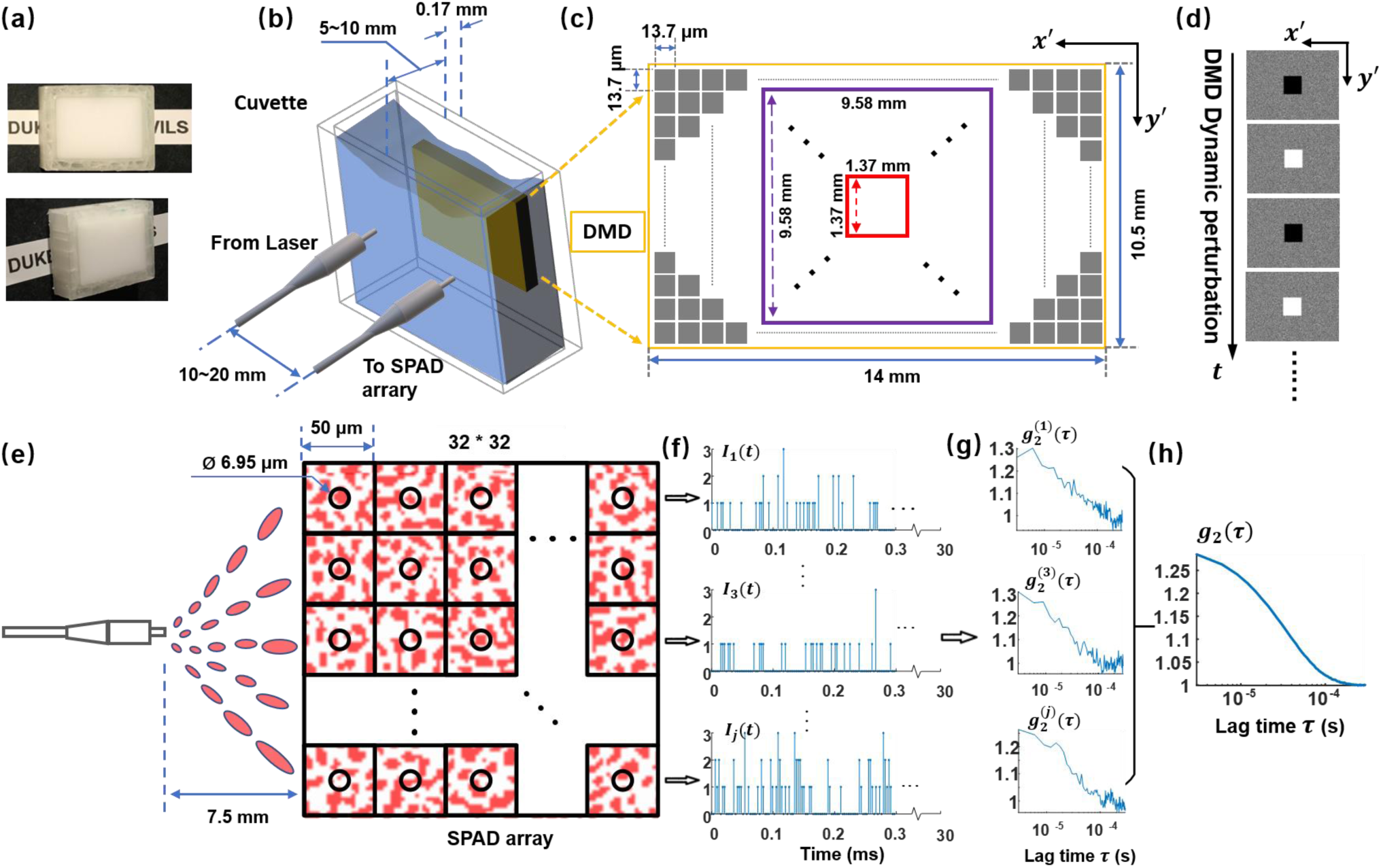
Detailed diagram of our parallelized DCS phantom setup for sensitivity and temporal resolution characterization. **(a)** Images of tissue phantom consisting of polystyrene microsphere-in-water suspension in a customized cuvette; **(b)** DMD phantom setup with DMD placed immediately behind tissue phantom. The distance between the centers of the illumination fiber and the detection fiber *d* was set to be approximately twice the cuvette thickness; **(c)** DMD panel. Square colored boxes represent different perturbation areas, varying from 1.37 × 1.37 mm ^2^ to 9.58 × 9.58 mm ^2^; **(d)** Example dynamic variation patterns, where pixels in middle square area switch between ‘on’ and ‘off’ states at a rate of multiple kHz (corresponding to the square color box in (c)), while the peripheral area is a constant random pattern; **(e)** Schematic layout of the 32 × 32 SPAD array illustrated with representative detected speckle patterns. The average speckle size at the SPAD array was tuned by adjusting detection fiber distance to approximately match the SPAD pixel active area (black circle). **(f)** Representative raw intensity measurements, *I*_*j*_(*t*), from each SPAD pixel with a temporal resolution of 3 μs. **(g)** Corresponding intensity autocorrelation curves of each SPAD pixel, 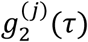, integrated over 30 ms, and **(h)** The final averaged autocorrelation curve *g*_2_(*τ*) using all 1024 SPADs.

## Results

### Phantom Setup

To provide a preliminary assessment of highly parallelized DCS based upon an integrated SPAD array, we constructed an easily reconfigurable phantom setup to create realistic kHz-rate perturbations deep within a decorrelating scattering slab. To achieve this goal, we turned to a non-biological phantom arrangement that offered a useful testing platform to quantitatively evaluate our new system’s performance gains. Multiple tissue-mimicking phantoms have previously been developed to simulate the dynamic scattering and absorption properties of human tissue such as that from the brain and skin^46-51^ (made of intralipid^48^, glycerol with fat emulsion^47^, or silicon rubber with small embedded scatterers^49,51^, for example). The dynamic optical properties of these prior phantoms have been typically tuned by varying the phantom material’s temperature, viscosity, or with pump-induced motion^47,50,52^. Since it is challenging to precisely vary the size and decorrelation rate of a hidden perturbation within these prior phantom setups, we instead created a digitally addressable phantom model that utilizes a DMD as the experimental perturbation source, which was placed immediately behind a thick and rapid decorrelating tissue phantom (5-10 mm thickness) with scattering properties that closely match living tissue (Fig.2 (b)). We varied the relative strength and speed of perturbations by changing both the size and frequency of projected patterns within a small DMD area (Fig.2 (c)), which provided a flexible means to mimic the effects of additional decorrelation induced by blood flow (for example). Measured decorrelation times arising from our tissue phantom generally fell between 10^−4^ and 10^−5^ s, with DMD-induced perturbations possible up to 22 kHz. We note that the DMD does not produce perturbations equivalent to any particular biological event. We instead used it here to facilitate quantitative analysis of the relative performance variations of parallelized DCS detection. In Supplementary Note 2, we also show that, with standard simulation tools^53^, the generated response of our DMD perturbations can be approximately related to the expected size of a biologically realistic inclusion embedded within a semi-infinite medium^54-57^. The schematic diagram of the DCS system for our DMD phantom study is shown in Fig. 1, and the setup parameters can be found in the methods section.

To maximize measurement SNR, it is beneficial to detect one to several optical modes (i.e., speckle grains) per SPAD^58^. To enable this criterion across the SPAD array, we ensured that the average speckle size contained within the optical field reaching the array is equal to, or a large fraction of, the size of each SPAD active area. This was achieved by adjusting the distance between the detection fiber and the photosensitive surface of the SPAD array (see Methods). We note that the relatively low fill factor of current SPAD array technology (7 μm SPAD active area with 50 μm pixel pitch in our setup, Fig. 2(e)) does not significantly impact parallelized DCS given the above measurement SNR constraint.

### Data processing

After detecting a sequence of SPAD array measurements, we first computed the autocorrelation curve at each SPAD pixel separately, and then averaged the computed autocorrelation curves into a final result (Fig. 2(f)-(h)). We note that there are many other possible methods to process the SPAD array’s raw 3D video frame data that we plan to examine in the future. To quantify the scattered field’s fluctuations, we computed the normalized temporal intensity autocorrelation function at the *j*th SPAD pixel, 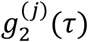, using the following equation,

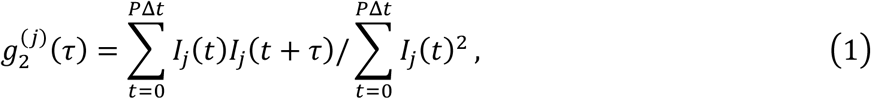

where *I*_*j*_(*t*) is the intensity of the *j*th SPAD in the array at a given time *t, τ* is the “lag” or correlation time, and we perform a summation over *P* temporal samples until reaching a desired degree of statistical averaging for one autocorrelation curve. In our phantom studies, each autocorrelation function *g*_2_ was typically calculated across *P* = 10,000 or 35000 temporal points, yielding one new autocorrelation curve every *P*Δ*t* = 30 ms or 105 ms, where Δ*t* = 3 μs is the SPAD’s temporal sampling rate. We will refer to *P*Δ*t* as the “autocorrelation curve rate”, which is different than the SPAD temporal sampling rate and defines the effective measurement rate for our system. After calculating 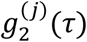 for each single SPAD (Fig. 2(g)), we then average these autocorrelations curves across *j* = 1 to *M* SPAD pixels to compute the final system autocorrelation function:

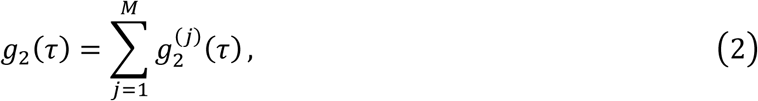

which yields a curve with higher SNR, as shown in Fig. 2(g-h). In most of our tests, we used a lag time that ranged from *τ*_*min*_ = 0 to *τ*_*max*_ = 300 μs for *g*_2_ computation, as we observed that curves decayed to approximately 1 when the lag time *τ* is larger than 200 μs.

The autocorrelation function, *g*_2_(*τ*), is related to the normalized electric field autocorrelation function, *g*_1_(*τ*), by the Siegert relation^59^:

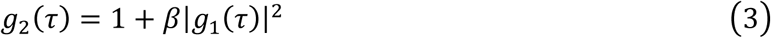

where *β* is a coefficient factor. To attain a decorrelation measure, we fit *g*_2_(*τ*) using the following equation,

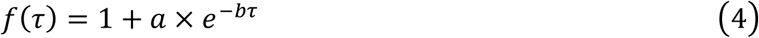

and report, *t*_*d*_, defined as the the delay time when *g*_2_ decays to 1/e of its initial value (*g*_2_(3*μs*)), as the “decorrelation time”. We sample *t*_*d*_ at the system autocorrelation curve rate *P*Δ*t*, and report the coefficient *b* of the index term as “decorrelation speed”.

### Increased accuracy and sensitivity of parallelized DCS using a SPAD array

Fig. 3 demonstrates the accuracy gained by using 1024 SPADs for parallelized DCS measurement. During this acquisition, we used a 5 mm thick tissue phantom with 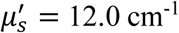 and did not induce any additional perturbation with the DMD, showing a tissue phantom decorrelation time of approximately 40 μs. Figure 3(a) shows representative raw data captured by a single SPAD over 10 ms. The average number of photons per pixel per sample is approximately 0.58. Figure 3(b)-(c) show the resulting autocorrelation curves generated at 10 ms and 100 ms autocorrelation curve rates. The blue curves represent the results using measurements from one SPAD, 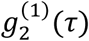, and approximately matches previous experimental measurements from common single-detector DCS setups. The red curves are the average of the autocorrelations *g*_2_(*τ*) from all *M* = 1024 SPADs within the array. The results show that with longer autocorrelation curve rates, the *g*_2_(*τ*) curve of a single SPAD smoothens to approximately match the curve produced by all 1024 SPADs, but when the autocorrelation curve rate is short, the results of a single SPAD not only have a low SNR, but also deviate from the averaged curve created by our parallelized approach.

**Fig. 3.**
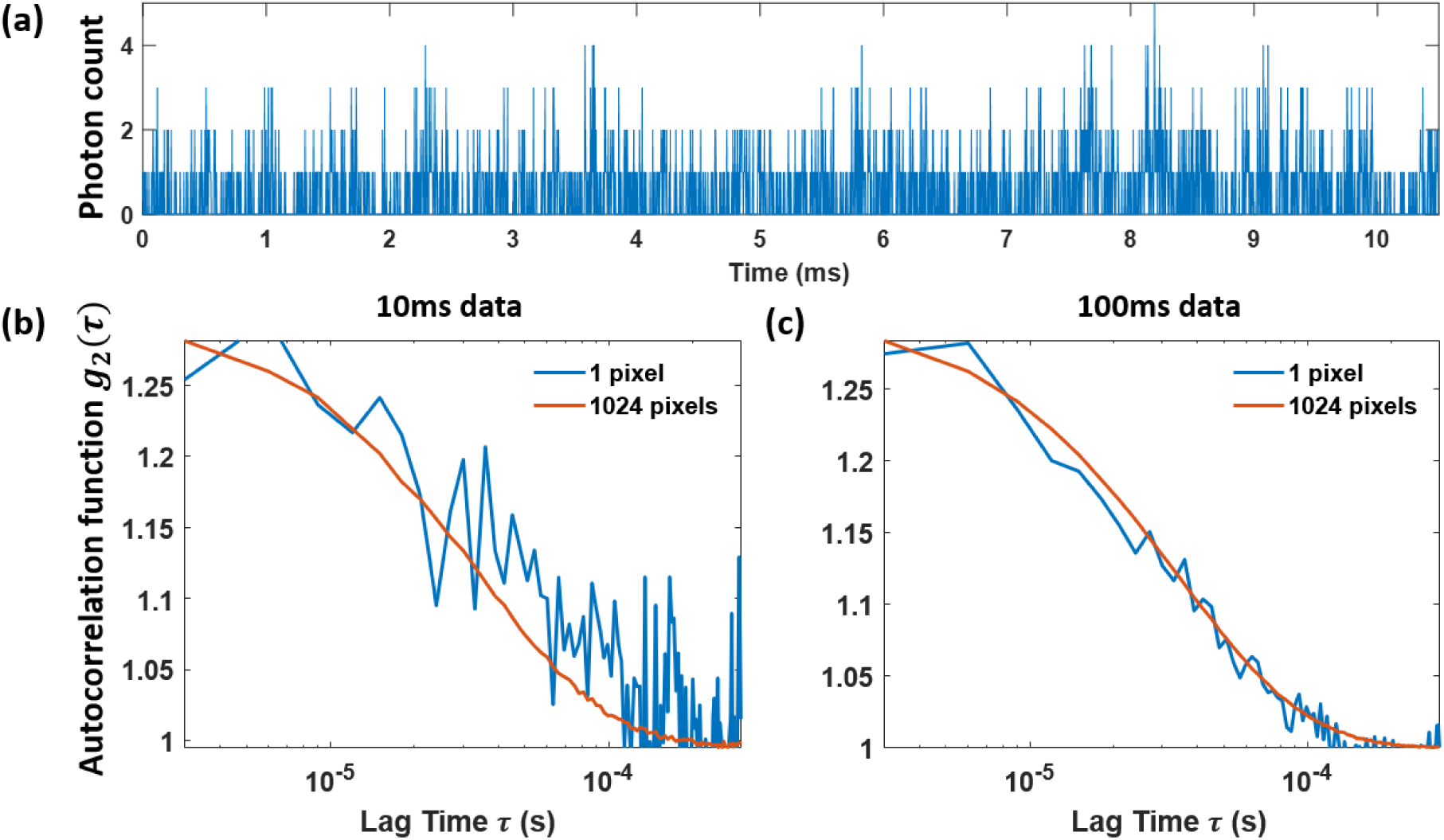
The average decorrelation curve generated by multiple SPAD array pixels increases DCS accuracy. **(a)** Raw data of temporal light intensity fluctuation from one SPAD pixel. Autocorrelation curves calculated using a single SPAD (blue) and all 1024 SPAD pixels (red) with a measurement window (i.e., autocorrelation curve rate) of **(b)** 10 ms and **(c)** 100 ms, respectively.

Next, to demonstrate the sensitivity gains of highly parallelized DCS, we tested the system’s ability to detect an embedded perturbation using a variable number of SPADs to compute a final *g*_2_(*τ*) curve. We compared the averaged *g*_2_(*τ*) curve that resulted from using 1, 9 and 1024 SPADs to detect a 4.1× 4.1 mm^2^ perturbation area, presented at 5 kHz, behind a 5 mm thick tissue phantom with 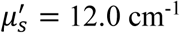. We first calculated the autocorrelation curve of each SPAD pixel using an autocorrelation curve rate of 30 ms (*P* = 10,000 temporal measurements detected at 333 kHz), and then averaged the resulting *g*_2_ curves from *M* = 1, 9 and 1024 SPADs, as shown in Fig. 4(a). After repeating this experiment over 420 independent trials, we computed the autocorrelation curve mean ± standard deviation (SD), displayed as solid center lines with a band of finite width in Fig. 4(b). The red and blue bands represent measurements performed with and without the perturbation present, respectively, where the measurement without a perturbation had the entire DMD area set to a random constant pattern. Here, we see that the averaged autocorrelation *g*_2_ curves are either completely indistinguishable or only partially distinguished when using either 1 or 9 SPADs, but become completely distinguishable (i.e., fully separated) when using 1024 SPADs, clearly reflecting an improved system sensitivity.

**Fig. 4.**
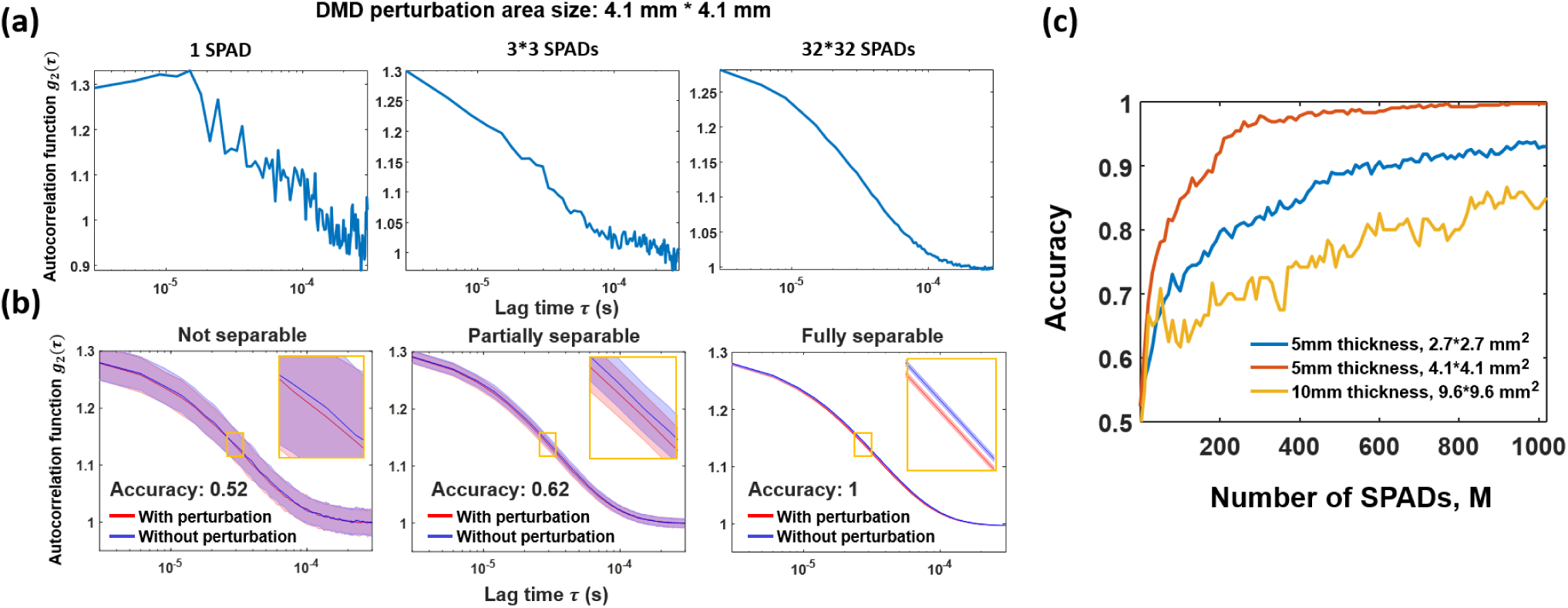
Demonstration of increased sensitivity to embedded perturbations with an increased number of SPADs. **(a)** Autocorrelation curves calculated using 1, 9 (3 × 3) and all 1024 (32 × 32) SPADs respectively, with an autocorrelation curve rate (*P*Δ*t*) of 30 ms and a 4.1 × 4.1 mm^2^ DMD perturbation area; **(b)** Autocorrelation curve separability and corresponding perturbation detection accuracy using 1, 9 and 1024 SPADs. Banded curves represent the mean ± standard deviation (SD) of 420 averaged autocorrelations, where red curve is measurement with perturbation, and blue curve is measurement without perturbation; **(c)** Perturbation detection accuracy increases with a larger number of SPADs for three example setup configurations: 5 mm phantom thickness and 2.7 × 2.7 mm^2^ perturbation area (blue), 5 mm phantom thickness and 4.1 × 4.1 mm^2^ perturbation area (red), and 10 mm phantom thickness and 9.6 × 9.6 mm^2^ perturbation area (yellow).

To quantify detection sensitivity, we computed a simple classification accuracy metric by alternately measuring the decorrelation time *t*_*d*_ with and without a DMD-induced perturbation and comparing the two adjacent measurements. If the *t*_*d*_ determined with the perturbation present was smaller than the *t*_*d*_ without the perturbation present, we assumed successful perturbation detection, and if not, we assume the detection is failed. This simple classification accuracy metric, defined as the proportion of successful detections over 420 trials, is plotted as a function of the number of SPADs (*M*) used to compute the average decorrelation curve (Fig. 4(c)). The classification accuracy generally increases with an increase in the number of SPADs. Different color curves represent different tissue phantom properties and DMD perturbation areas and the accuracy increases when using a thinner tissue phantom or larger perturbation area.

### Minimally detectable DMD perturbation area

For a specific tissue phantom, the ability to detect a perturbation depends on both the perturbation area size and autocorrelation curve rate. We thus varied both of these parameters while repeating our classification accuracy tests. The results of this exercise are summarized in Figs. 5-6. For each setting, we repeated 120 independent experiments to calculate the classification accuracy. The mean ± SD of several example *g*_2_ curves (5 mm thick tissue phantom with 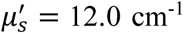) are in Fig. 5, where once again red and blue curves represent the case of perturbation present versus absent, respectively. The curves within each row use the same autocorrelation curve rate, while the curves within each column have the same DMD perturbation area. The results show that when using an autocorrelation curve rate of 10 ms, it is possible to accurately identify a 3.42 × 3.42 mm^2^ perturbation via the *g*_2_ curve separation. When the autocorrelation curve rate increases to 30 ms or 100 ms, it is possible to detect smaller perturbations, down to sizes of 2.74 × 2.74 mm^2^ and 2.05 × 2.05 mm^2^, respectively.

**Fig. 5.**
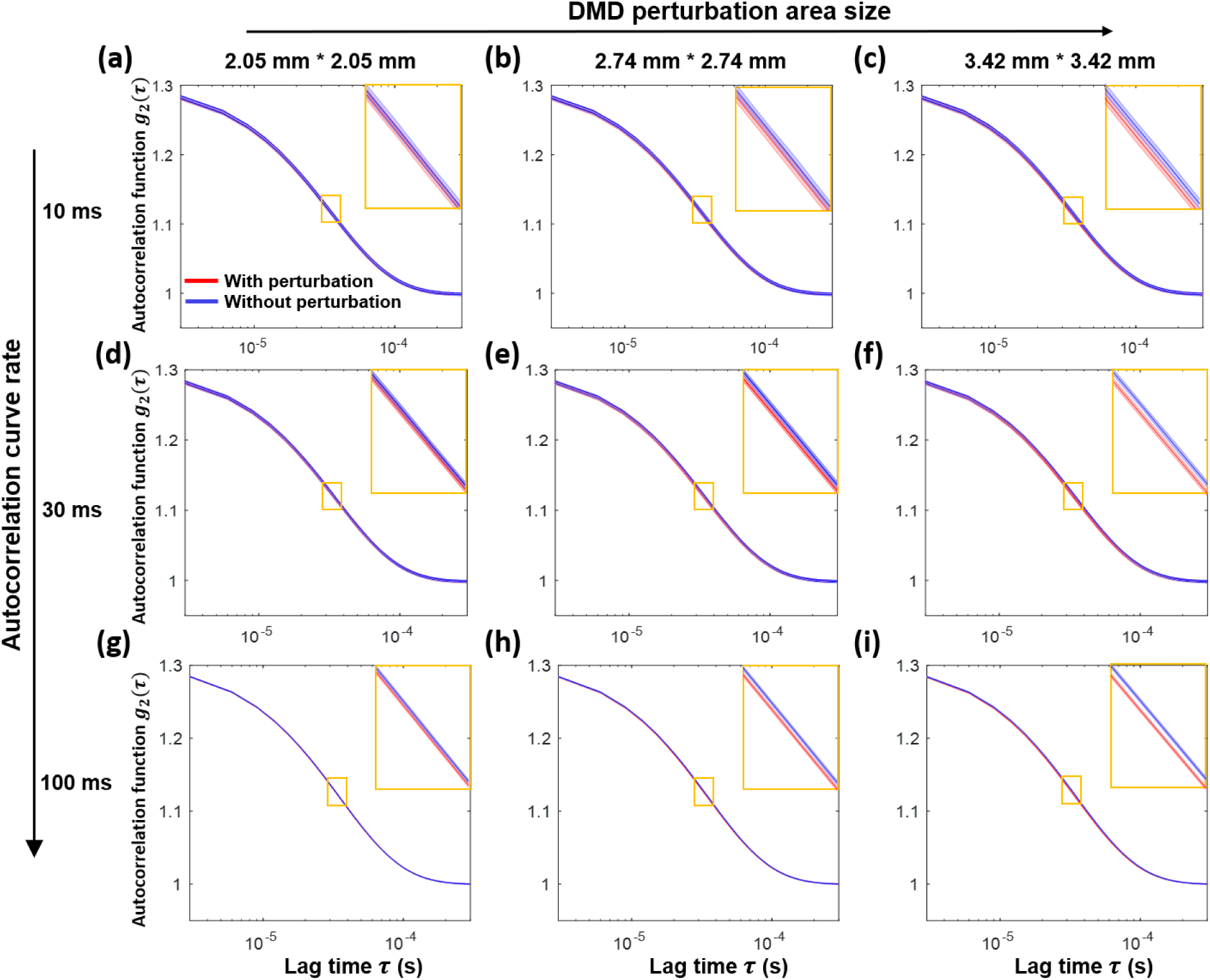
Parallelized DCS detection of mm-scale size perturbations at different autocorrelation curve rates. **(a) - (c)** Detection results with 10 ms autocorrelation curve rate for perturbation areas of size 2.05 × 2.05 mm^2^, 2.74 × 2.74 mm^2^ and 3.42 × 3.42 mm^2^, respectively. Banded curves represent the mean ± standard deviation (SD) of 120 averaged autocorrelation curves, where the red curve indicates with perturbation, blue curve band indicates without, and orange box is a selected zoom-in. **(d) - (f)** Similar detection results using a 30 ms and **(g) - (i)** 100 ms autocorrelation curve rate.

**Fig. 6.**
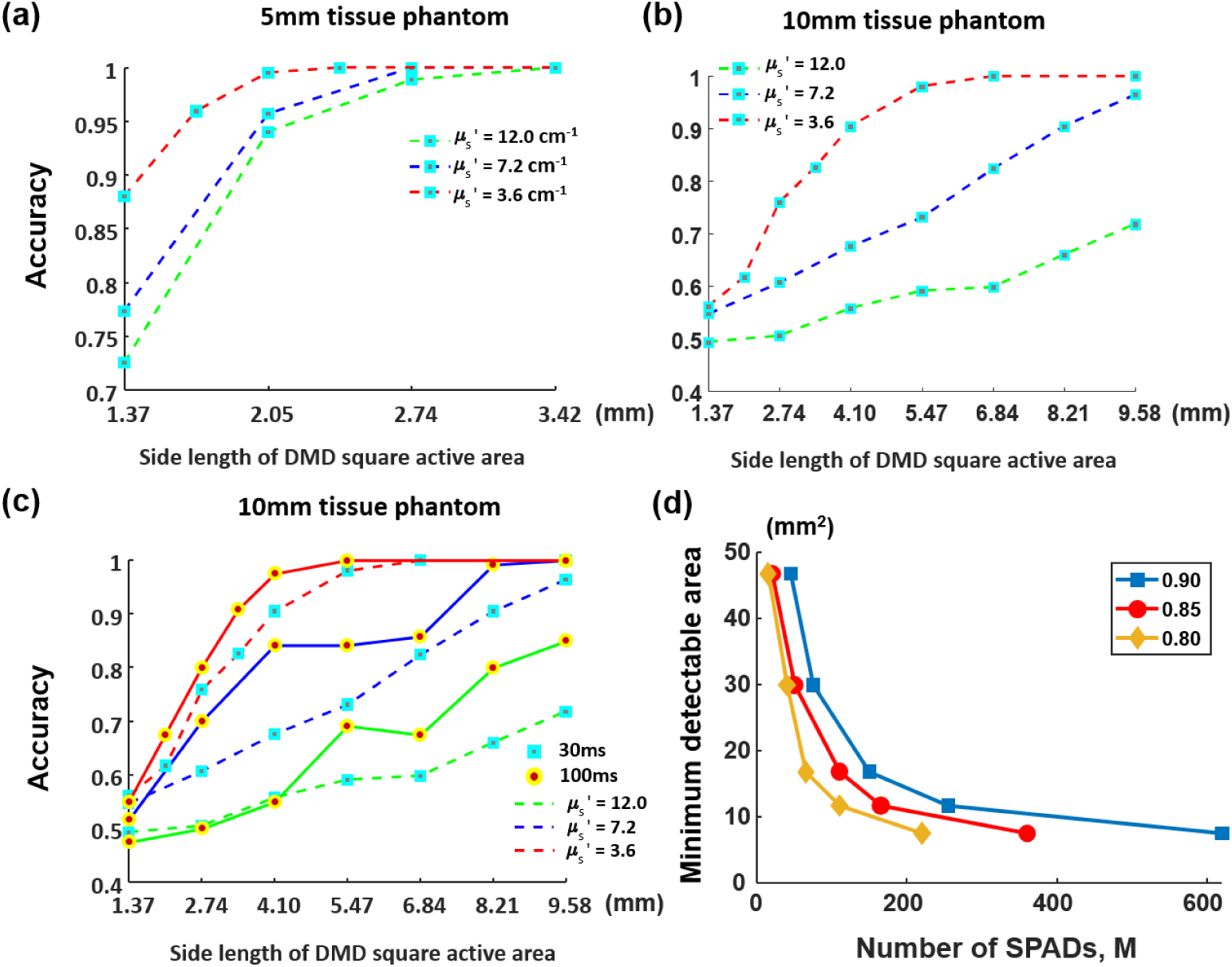
Parallelized DCS classification accuracy of different perturbation sizes, tissue phantom properties and autocorrelation curve rates. **(a)** Classification accuracy increases for larger perturbation areas and lower reduced scattering coefficients. Here, 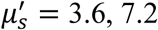, and 12.0 cm^-1^ for red, blue and green curves respectively, for 5 mm tissue phantom thickness and 30 ms autocorrelation curve rate. (b) Matching results with 10 mm thick tissue phantom. (c) Comparison of classification accuracy of different autocorrelation curve rates with 10 mm tissue phantom. Dotted line is for 30 ms and solid line is for 100 ms. (d) The minimum detectable perturbation area as a function of the number of SPADs. Yellow, red and blue curves represent results corresponding to classification accuracy thresholds at 0.8, 0.85 and 0.9 for 5 mm tissue phantom thickness and 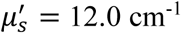.

Figure 6(a)-(b) summarizes the classification accuracy of our parallelized DCS system under different phantom scattering coefficients 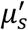 and for different perturbation sizes, for a 5 and 10 mm thick tissue phantom, respectively. A 30 ms autocorrelation curve rate was used to calculate *g*_2_ in (a) and (b). It is clear that the classification accuracy increases with a larger perturbation area and a lower tissue phantom reduced scattering coefficient and thickness. Classification accuracy also improves with a longer autocorrelation curve rate (100 ms vs. 30 ms, Figure 6(c)). Figure 6(d) plots the minimum detectable perturbation area at different detection accuracy thresholds as a function of number of SPADs *M* used for parallelized detection (5 mm thickness, 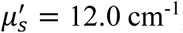, autocorrelation curve rate of 30 ms). These results show the significant improvement in system sensitivity with increased measurement parallelization.

### Impact of perturbation frequency on autocorrelation function

To test the effect of the temporal perturbation rate on the average autocorrelation function of our parallelized DCS system, we varied the refresh frequency of the DMD perturbation area between 1 kHz and 20 kHz for 3 different perturbation area sizes, with the results shown in Fig. 7. The tissue phantom thickness in this experiment was 5 mm with 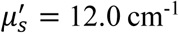. With an autocorrelation curve rate of 30 ms, the *g*_2_ curve clearly dips with increasing DMD frequency, and it is easier to differentiate *g*_2_ curves corresponding to different perturbation frequencies for larger perturbation areas, as expected.

**Fig. 7.**
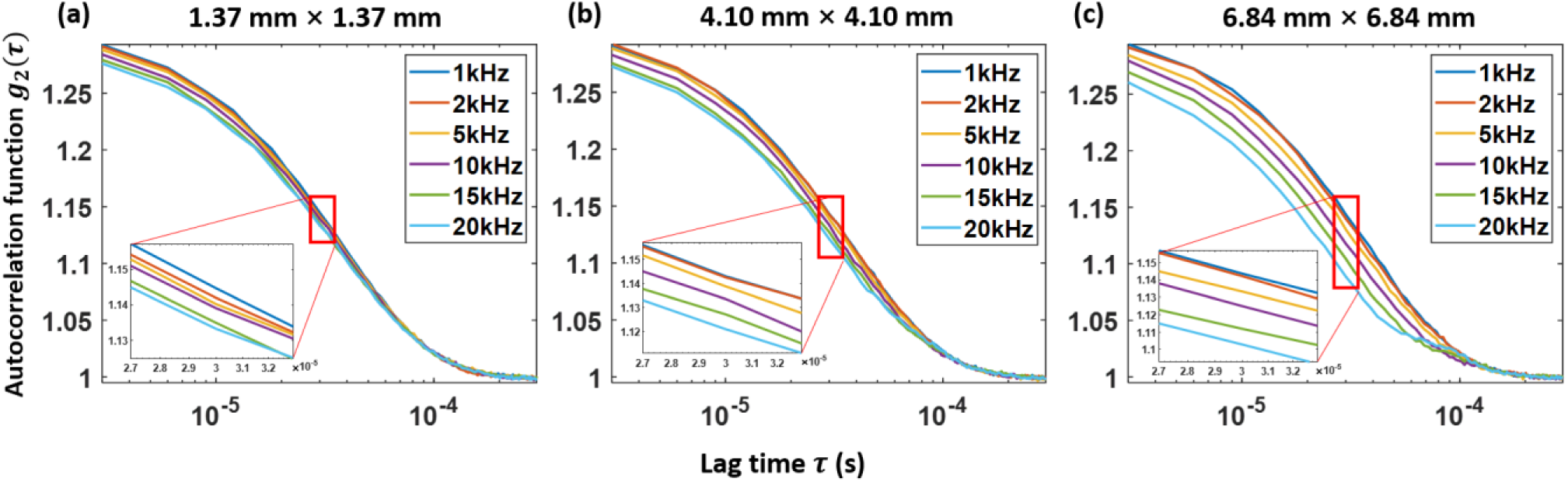
The effect of perturbation’s temporal rate on the average autocorrelation function of the parallelized DCS system. Different colored autocorrelation curves represent different perturbation frequencies. The perturbation areas corresponding to (a), (b) and (c) are: 1.37 × 1.37 mm^2^, 4.10 × 4.10 mm^2^ and 6.84 × 6.84 mm^2^, respectively. The autocorrelation curve rate is 30 ms.

### *In vivo* forehead blood flow results

To demonstrate that our highly parallelized DCS system can also improve the SNR of *in vivo* measurements, we modified our setup to measure blood flow within the human forehead and verified our measurements with a commercial pulse oximeter. The source-detector distance *d* was set at 1.5 cm and each measurement consisted of 21 s of continuous frame data. We processed the data to generate decorrelation speed plots sampled at different autocorrelation curve rates, *P*Δ*t*, as well as with using measurements from a variable number of SPADs, *M*, to demonstrate the temporal resolution advantages of our parallelized approach. Example results from these tests are shown in Fig. 8. Similar to our phantom studies, we fit the autocorrelation curves using an exponential function and extract and plot the decay coefficient *b* (Eq. (4)). A larger *b* value represents faster decorrelation speed (i.e., faster blood flow).

**Fig. 8.**
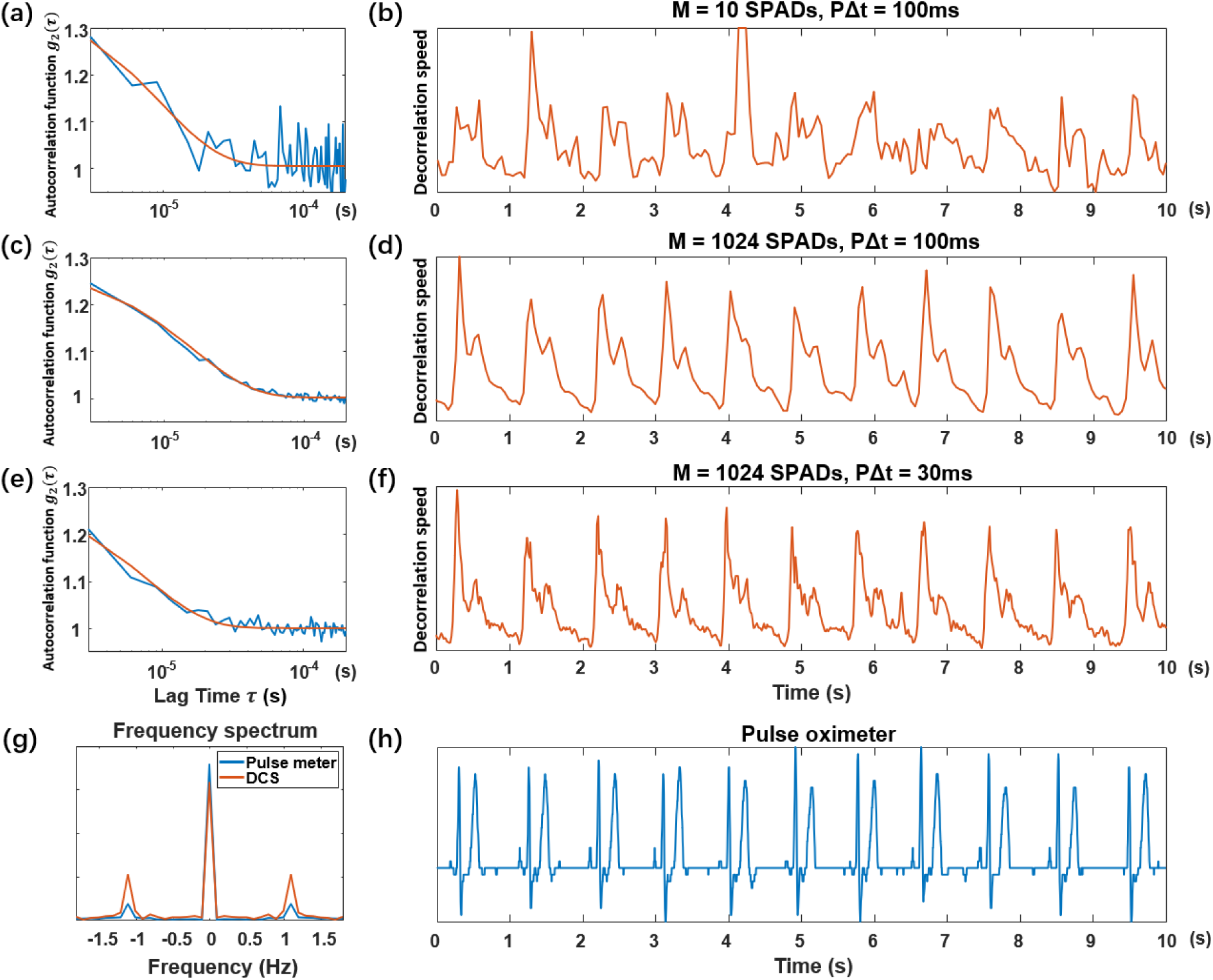
Human forehead pulse measurement results. Representative autocorrelation curves (blue) and their best exponential function fit (red) are at left (a, c, e). At right are associated DCS pulse measurements over 10 s (b, d, f), which are represented by the plot of normalized decorrelation speed, extracted from autocorrelation curves after exponential fitting. (a-b) Using *M* = 10 SPADs and an autocorrelation curve rate *P*Δ*t* = 100 ms, (c-d) *M* = 1024 SPADs and an autocorrelation curve rate of *P*Δ*t* = 100 ms, and (e-f) *M* = 1024 SPADs and an autocorrelation curve rate, *P*Δ*t* = 30 ms. (g) The Fourier transform of the DCS pulse signal (f) and the pulse signal acquired using a commercial pulse meter (h).

The representative autocorrelation curves and the pulse signals calculated using the normalized decorrelation speed *b* were shown in Fig. 8(a-f). With an autocorrelation curve rate of *P*Δ*t* = 100ms and averaging all *M* = 1024 SPADs, we obtain a high-SNR autocorrelation function *g*_2_(*τ*) (Fig. 8(c)) with a decorrelation time that clearly follows the periodicity of the subject pulse (Fig. 8(d)). Two deflections caused by ventricle contractions and repolarization can be clearly observed. However, if only *M* = 10 SPADs were used for averaging, pulse signal retrieval remains challenging (Fig. 8(b)), as the decorrelation speed *b* extracted from the exponential fit is heavily affected by a low SNR autocorrelation function *g*_2_(*τ*) (Fig. 8(a)). Moreover, we are still able to retrieve the pulse signal with a shorter autocorrelation curve rate, *P*Δ*t* = 30 ms (Fig. 8(e-f)). This is further validated by examining the Fourier transform of the DCS decorrelation speed signal (Fig. 8(f)) and the pulse signal simultaneously acquired by the pulse oximeter (Fig. 8(h)), shown in Fig. 8(g), which have approximately the same peak frequency at ∼1.1 Hz. Consistent results were obtained for 5 different human subjects.

### Human prefrontal cortex activation test

As an additional demonstration of the sensitivity gains of highly parallelized DCS, we also measured dynamic variations induced by a behavioral task. The prefrontal cortex region is considered to manage in part cognitive processes such as planning, cognitive flexibility, and working memory^60^. Previous studies with the complementary non-invasive optical measurement approach termed interferometric near-infrared spectroscopy (iNIRS), for example, measured and reported dynamic changes within the prefrontal cortex area for subjects who read a paragraph of unfamiliar text after a 10-min rest period^6^. Here, we explored the likelihood of observing such dynamic activity using autocorrelation functions obtained from our highly parallelized DCS system. The subjects first rested for 5 minutes, and then began a 15-minute test divided into three stages: reading stage 1, resting stage, reading stage 2, where each stage lasted 5 minutes. We collected DCS measurements for the first 10 seconds of every test minute using all 1024 SPADs. The autocorrelation function *g*_2_(*τ*) and the associated decorrelation time *t*_*d*_ were calculated using an autocorrelation curve rate of 100 ms. A representative plot of the decorrelation time measurement of one subject is shown in Fig. 9(a). We then calculated the mean decorrelation time for each subject by averaging the 100 values of *t*_*d*_ that comprise each 10 s DCS measurement window (see plot in Fig. 9(b)). A total of 6 subjects were recruited and tested in this study. To first detect and remove subject data that included unwanted motion artifacts during each-minute test, we calculated the standard deviation (SD), σ_d_, of the decorrelation time, t_d_, within each 10s measurement window (15 in total, Supplementary Fig. 3) to obtain the 95% confidence interval of σ_d_ (±2σ_d,_ red dash lines in Supplementary Fig.3, 90 total measurements), and then removed experiments exhibiting a decorrelation time standard deviation σ_d_ outside of the confidence interval. In this study, subject #3 and #6 included such data, suggesting significant motion occurred, which we aim to address with a new setup in future work (see Discussion). The mean decorrelation time of the remaining 4 subjects are shown in Fig. 9(c) with the mean ± SD shown as a dotted line with a banded curve. The solid horizontal line represents the mean decorrelation time across each of the three test stages (the first reading stage, resting stage, and the second reading stage). The mean decorrelation time in both reading stages (blue) was generally observed to be lower than in the resting stage (red), indicating an increase in blood flow speed within the measurement area (i.e., activation of prefrontal cortex), matching prior results^6^.

**Fig. 9.**
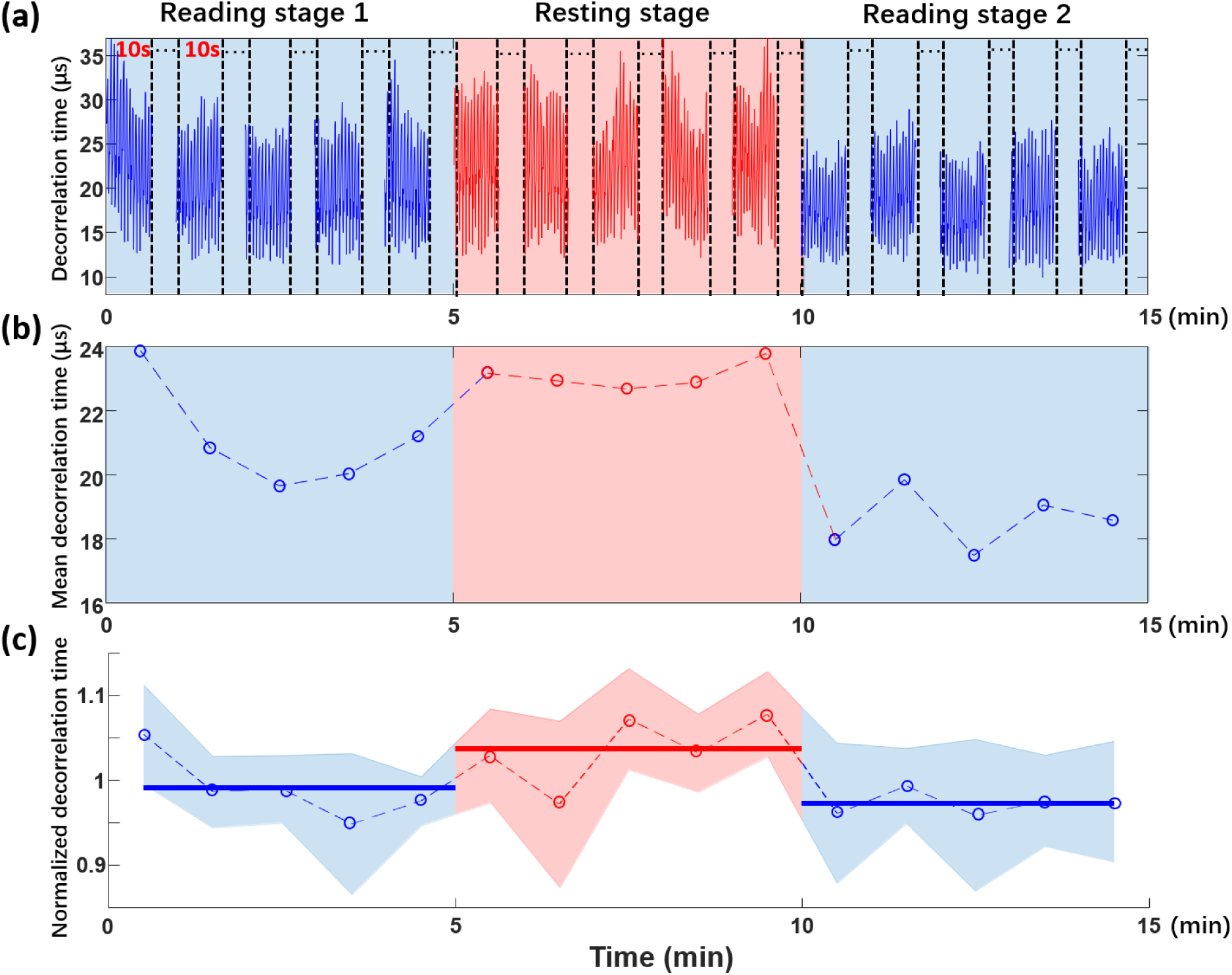
Human prefrontal cortex activation measurement. **(a)** Plot of the decorrelation time (*t*_*d*_) values over 15 minute test including two reading stages and one intermediate rest stage, were 10 s of signal is collected every minute. **(b)** Plot of mean decorrelation time corresponding to (a). Each mean decorrelation time value is obtained by averaging all the decorrelation time values within corresponding 10 s window. **(c)** Mean ± SD results of the mean decorrelation time of 4 subjects. The decorrelation time of each subject is obtained after normalizing their mean value to 1. Solid horizontal lines represent the average of the five normalized decorrelation times in each stage.

## Discussion

In this study, we demonstrated the highly parallelized acquisition of DCS signal using an integrated SPAD array that contained 32 × 32 pixels, which allowed us to average over a thousand independent speckle fluctuation measurements per frame to improve system sensitivity and speed. We found that the SPAD array provides enhanced sensitivity as compared to a single or a small number of SPADs by testing its performance with a novel perturbation model, implemented by inserting a DMD beneath a thick tissue phantom, which demonstrated accurate detection of mm-scale perturbations under 1 cm of tissue-like phantom. We also used this new approach to measure the blood pulse from the human forehead with a 30-millisecond temporal resolution and detect behavior-induced variations from above the pre-frontal cortex.

Monolithic CMOS SPAD arrays continue to rapidly increase in pixel count (e.g., an 1 megapixel array is now available^61^), and recent consumer-oriented developments have led to combined sensing and processing layers via 3D stacking that open up new functionalities^33^. Like other integrated CMOS technology, we expect both pixel counts and functionality to increase with improvements from an ever-advancing semiconductor manufacturing industry. Prior work has shown that more than one single-photon detector can increase DCS measurement sensitivity for the challenging task of non-invasive detection of functional brain dynamics^13,62^, and tomographically measure cerebral blood flow^50,63^, for example. Our new approach here offers a means to implement such efforts with dozens of times more detectors and thus increase speed or sensitivity by a proportional factor, thus opening up a new suite of possible experiments.

The sampling rate of our SPAD array in this study is 333 kHz, which is slower than the typical sampling rate (1 MHz or more) of single SPADs in previously reported DCS setups. Nevertheless, as our *in vivo* study and phantom-based experiments demonstrate, this lower sampling rate is still fast enough to retrieve the entire autocorrelation function of relevant biological activity such as blood flow with a source-detector distance of 1.5 cm. Our SPAD array currently operates at a bit depth of 8, which is more than sufficient for most of DCS applications. In future work, we aim to work at an increased sampling rate of 720 kHz with a decreased per-sample bit depth, which is possible with our current hardware. In addition, we used a 670 nm illumination source, as the detection sensitivity of our SPAD array is higher in the visible wavelength range. However, a longer wavelength would potentially increase the penetration depth into deep tissue. Future studies will thus aim to further optimize the selected illumination wavelength for a particular task, and to optimize our human subject setup to minimize the impact of slight head movements. Although we adopted several measures to keep the subject’s head as stable as possible (see Methods), we occasionally observed dramatic changes in the mean decorrelation time value during long-term measurement, which may have been caused by the changes of the multimode fiber’s position and/or pressure on the skin surface. To improve setup stability, we aim to modify our headwear setup and design appropriate behavioral tasks that can be completed by reclining participants. Finally, the large spatio-temporal datasets acquired per experiment suggest that a new suite of software tools are on the horizon for highly parallelized DCS setups.

## Methods

### Phantom experimental setup

Our decorrelating tissue phantom was created by a 1 μm polystyrene microsphere-in-water suspension (Polysciences, inc.; Warrington, PA) in custom made cuvettes. The front and back walls of the cuvette perpendicular to the fiber tips were made of microscope coverslips with a thickness of 0.17 mm. The back wall coverslip was placed directly against the surface of a DMD (Digital Light Innovations; Austin, TX; 1024 × 768 micromirrors with a micromirror pitch size of 13.68 μm) and completely covered its surface (see Fig. 2(c)). In this study, we tested two different phantom tissue thicknesses using cuvettes with thicknesses of 5 mm and 10 mm (Fig. 2(b)). We also tested scattering tissue phantoms with three different reduced scattering coefficients: 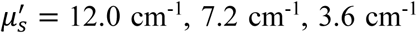, created by varying the microsphere-in-water concentration. These coefficient values are close to previously reported reduced scattering coefficients of human skin, brain, and other soft tissue^64^.

The DMD could generate binary patterns at a frequency of up to 22 kHz. The middle square area of the DMD (the “perturbation area”) switches between the ‘on’ and ‘off’ states at a rate of multiple kHz to simulate changes caused by deep tissue motion, while the peripheral area is a random pattern that remains constant. We varied the size of the central perturbation area from 1.37 × 1.37 mm^2^ (100 × 100 micromirrors) to 9.58 × 9.58 mm^2^ (700 × 700 micromirrors), as shown in Fig. 2(f). The measurement without the DMD-induced perturbation is achieved by setting the entire DMD area to be a random pattern that remained constant.

Light from a 670 nm long coherence laser (Opto Engine LLC; Midvale, UT) with 150 mW input power was delivered by a multimode fiber(core diameter = 50 μm, NA = 0.22, Thorlabs; Newton, NJ) to the surface of the tissue phantom. To match power conditions required during *in vivo* experiments, we also followed the American National Standard for Safe Use of Lasers (ANSI) power limit for our phantom studies, which requires an irradiance less than 200 mW/cm^2^. We placed the illumination fiber tip 22 mm away from the cuvette surface to form a 10 mm diameter spot on the phantom surface, which provided our system with a peak irradiance of about 191 mW/cm^2^. We set the distance between the centers of the illumination fiber and the detection fiber (core diameter = 1000 μm, NA = 0.5, Thorlabs; Newton, NJ) to be twice the thickness of the phantom to collect photons following the banana shaped path’s most probable trajectories (Fig. 2 (b))^1^. The SPAD array (Photon Force PF-32) consists of 32 × 32 single SPADs, and each single SPAD is 50 μm × 50 μm in size, with a circular active area of approximately 7 μm in diameter (Fig. 2(e)). The sampling rate of each SPAD is 333kHz with a bit depth of 8.

### Geometry for parallelized speckle detection with SPAD array

To tune the average speckle size at the detector plane, we placed the detector fiber tip at an appropriate distance (*R*) from the SPAD array. Selecting a distance between the fiber and the SPAD array that is too small will cause each SPAD to contain many independent speckle grains, which leads to a reduced dynamic speckle contrast, while selecting a distance that is too large will yield a low SNR. To match the average speckle size to the SPAD pixel active area, which yields maximum contrast, we calculated the speckle grain size *d*_*s*_ at 50% correlation generated from a fiber as a function of the distance and fiber core diameter using the following equation^65^:

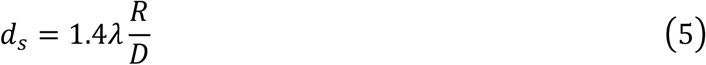

Here, *λ* is the wavelength of the illumination laser, *R* is the distance between the detection fiber and SPAD array, and *D* is the core diameter of the detection fiber. To obtain the maximum SNR, *d*_*s*_ should be similar to the diameter of the SPAD pixel active area of 7 μm. In our setup, we used a multimode fiber with *D* = 1 mm, making the ideal fiber-to-SPAD distance *R* = 7.5 mm.

### Stability of multiple tests in calculating classification accuracy

In the tissue phantom setup, motion of both the DMD and microsphere-based tissue phantom potentially affect the SPAD array signal. During our experiments, it was important to keep the microsphere solution stable in terms of concentration and temperature to maintain a constant phantom-induced decorrelation rate. To achieve this, we placed the entire system in a closed box to reduce the effect of air flow on the solution, and illuminated the sample and ran the DMD at the experimental frequency for 30 minutes before each experiment, to achieve a stable temperature and microsphere phantom movement before initiating data collection. To confirm this stability, we would collect a 3-minute measurement with the same DMD projection pattern, calculate a continuous autocorrelation function *g*_2_ with an autocorrelation curve rate of 0.1 s using Eq. 2, and ensure the decorrelation curves remained unchanged within the expected standard deviation.

### Human forehead blood flow and prefrontal cortex activation measurement setup

Similar to the phantom study setup, the laser (670 nm) was delivered by the same multimode fiber to the participant’s forehead. We adjusted the laser input power to 40 mW and attached a 3d printed holder to the fiber tip. The holder kept the fiber head ∼1.1 cm away from the subject’s forehead so that the illumination spot on the skin surface spread to 0.2 mm^2^, which ensured power irradiance below the ANSI limit (200 mW/cm^2^). Contact between the holder and forehead also helped reduce motion effects. The source-detector distance *d* was set at 1.5 cm and the SPAD array sampling rate was 333kHz. The participant’s head was rested on a chinrest and secured with an adjustable elastic bandage to further reduce the motion effects during the measurement, especially for the prefrontal cortex activation measurement. The participants also wore laser safety goggles (LG4, Thorlabs; Newton, NJ) during the whole experiment for eye safety. All the *in vivo* studies were approved by the Duke Campus Institutional Review Board.

For the initial forehead blood flow validation study, DCS measurements were taken for 21 seconds. To validate the accuracy of our DCS pulse measurement, a commercial pulse oximeter (EMAY Ltd; HongKong, China) was used to record the pulse signal simultaneously. For the prefrontal cortex activation measurement, we designed a test lasting for 20 minutes that was divided into four stages: 1) close eyes and rest for 5 minutes, 2) read for 5 minutes (reading stage 1), 3) close eyes and rested for 5 minutes, 4) reading for another 5 minutes (reading stage 2). During the test, we kept the environment dark and quiet, placed a tablet in front of the subjects that displayed preselected novels that the subjects had never read before. At the beginning of each stage, a beep sound was played to inform the subject to open eyes and start to read or close eyes and rest. Data transfer limitations from the SPAD array to computer memory limited the streaming data duration to 10 s per minute across all 1024 SPADs. The decorrelation curve rate for this experiment was 100 ms.

## Data availability

All relevant data are available from the authors upon request.

## Code availability

All relevant codes are available from the authors upon request.

## Acknowledgments

Research reported in this publication was supported by the National Institute of Neurological Disorders and Stroke of the National Institutes of Health under award number RF1NS113287, as well as the Duke-Coulter Translational Partnership. W.L. acknowledges the support from the China Scholarship Council. The authors would like to thank Kernel Inc. for their kind assistance.

## Author contributions

R.H. conceived and supervised the project. W.L. and R.Q. built the setup. W.L., R.Q. and S.X. programmed the reconstruction algorithm and analyzed the data. W.L. performed the phantom experiments. W.L., R.Q. and S.X. performed the *in vivo* experiments. S.X., E.B., J.J. performed the numerical simulations. W.L., R.Q., P.K., M.H., R.H., H.W., and Q.D. discussed the experimental implementation. W.L., R.Q., S.X., P.K., R.H., D.B. and C.C. discussed the results. W.L., R.Q., S.X. and R.H. wrote the manuscript with input from all the authors.

## Additional information

