## Supplementary Information for "Fast and sensitive diffuse correlation spectroscopy with highly parallelized single photon detection"

### Supplementary Note 1: Simulation of photon flux at tissue surface

In this section, we present details about a simulation to establish the average number of photons per speckle per second available at the surface of a tissue-like scattering slab in a DCS-type reflection configuration. These values were cited in the introduction to motivate the need for a parallelized detection strategy at large source-detector separations, which are desired to probe deep within scattering media. In this simulation, we use an open-access online Monte Carlo simulation tool to simulate light scattering through large scattering phantoms[1]. Three different phantoms materials were tested: Tissue I has $\mu_a = 0.4\ \mathrm{cm}^{-1}$ and $\mu_s' = 12.0\ \mathrm{cm}^{-1}$, Tissue II has $\mu_a = 0.2\ \mathrm{cm}^{-1}$ and $\mu_s' = 7.2\ \mathrm{cm}^{-1}$ and Tissue III has $\mu_a = 0.1\ \mathrm{cm}^{-1}$ and $\mu_s' = 3.6\ \mathrm{cm}^{-1}$. To match the experimental setup (see *Phantom experiment setup* in Methods section), we place the source 22 mm away from the surface, and diverged the incident beam to a circular spot with a diameter of 8 mm. 100 billion photons were sent into each tissue phantom, and the number of detected photons were then normalized to match the number of photons generated by a 100 mW continuous-wave laser beam at 670 nm, to meet ANSI safety requirements, using the Planck–Einstein relation.

Supplementary Figure 1 (a) illustrates the tissue phantom and illumination configuration used for the simulation study. Supplementary Figure 1(b) plots the number of detected photons per average speckle area per microsecond on the tissue surface at a given distance from the center of the incident source beam. The light transport inside the tissue phantom follows a banana-shaped path from the source to the detector, and spreads along the surface as depicted in Supplementary Figure 1(c). We assumed that the average area for a fully developed speckle is $(\lambda/2)^2$ [2].The number of detected photons at a source-detector separation of 14 and 21 mm, as used in our phantom experiments, are listed in supplementary Figure 1 (d). As is clear, the number of photons detected per speckle area at centimeter-scale distances from the incident source drops dramatically as the phantom's reduced scattering coefficient becomes larger (i.e., with more scattering). This low photon issue limits the sensitivity of DCS measurements and impacts the estimation of an accurate correlation curve, which is typically resolved by averaging the correlation curve over a longer sliding window, at the expense of decreasing the overall temporal resolution of each reported decorrelation value for the DCS system. With parallelized detection, as used in our experiments, averaging time periods can be shortened by orders of magnitude while still maintaining the same detection accuracy.

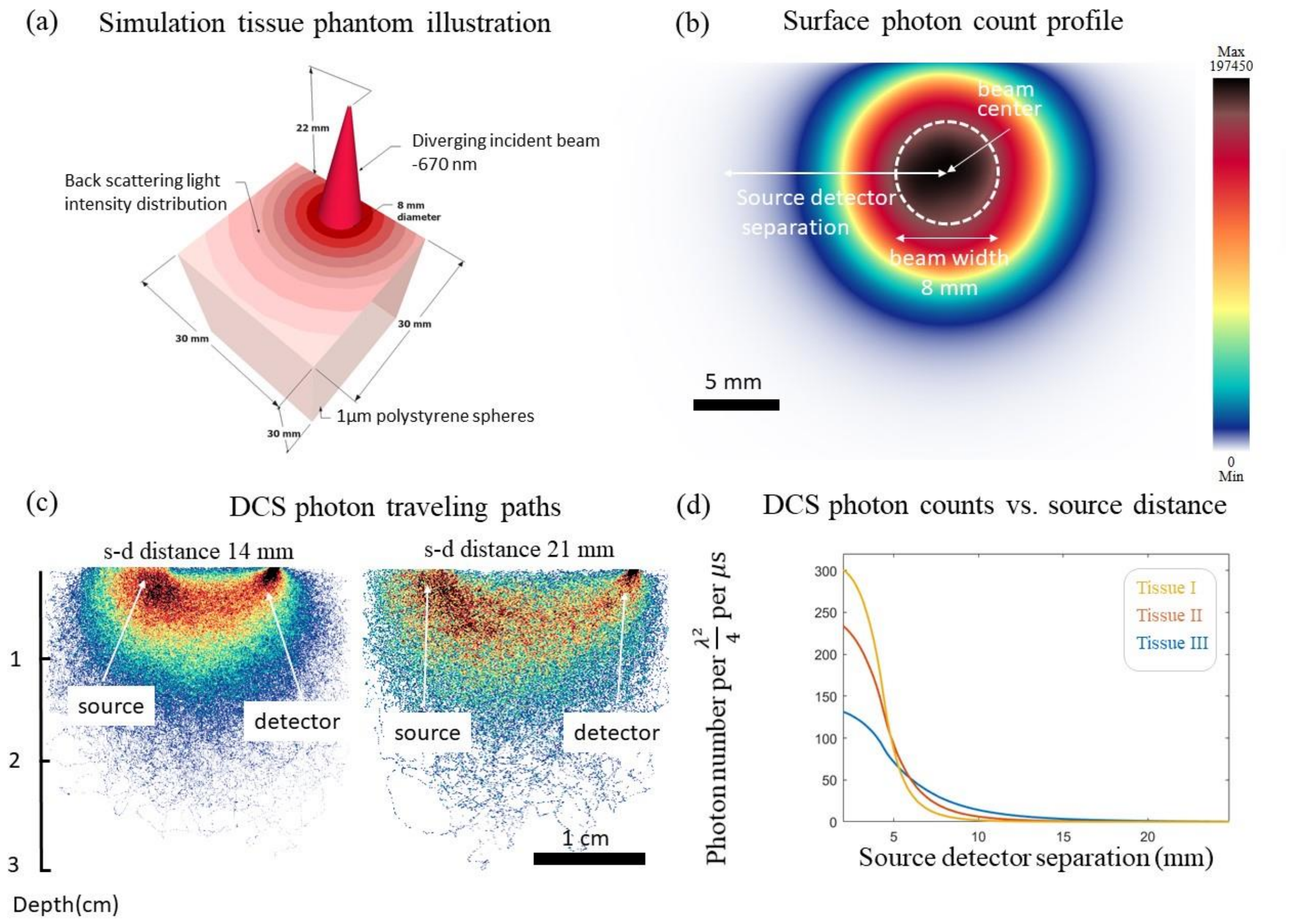


**Supplementary Figure 1. (a)** An illustration of the tissue phantom used for this simulation. The confined beam is 22 mm above the phantom and diverges to a 8 mm spot at the surface **(b)** Photon density map within the phantom tissue (tissue III) when 100 billion photons are used for illumination. **(c)** Simulated photon paths for source-detection separations of 14 and 21 mm. The banana-shaped photon path dips 3-10 mm and 6-15 mm for 14 mm and 21 mm source-detector separation, respectively **(d)** Plot of number of photons per averaged speckle area $(\lambda/2)^2$ per microsecond at different source-detector separations for three different types of homogeneous tissue phantom.

| | Tissue I | Tissue II | Tissue III |
|---|---|---|---|
| $\mu_s'$(mm$^{-1}$) | 1.20 | 0.72 | 0.36 |
| $\mu_a$(mm$^{-1}$) | 0.04 | 0.02 | 0.01 |
| # photon (14 mm) | 0.70 | 9.36 | 76.54 |
| # photon (21 mm) | 0.03 | 1.09 | 17.14 |

**Supplementary Table 2.** Detected photons per $\lambda^2/4$ per μs at 14 and 21 mm source-detector separations (as used in experiments) for three different phantom tissue types.

### Supplementary Note 2: Relation between DMD phantom perturbation experiment and blood flow model

One of the most widespread DCS applications is to measure blood flow (BF) beneath tissue[3,4]. To motivate the utility of our new DMD-based phantom setup for parallelized DCS system characterization, we attempt here to relate our experimental DCS measurements from this phantom arrangement to prior models and measurements of BF-induced DCS signal. For this task, we adopted a separate Monte Carlo simulation tool to track photon propagation and generate field and intensity autocorrelation curves from simulated blood flow[5].

In the following, we first present the mathematical setup of this DCS BF simulation, before showing results that allow us to connect BF measurements to our experimental results. Considering the $n^{th}$ photon experiencing its $i^{th}$ scattering inside the medium $m$, we can express the resulting momentum transfer at each scattering event as $\mathbf{q}_{n,m}^{i}$ and the photon travel path length between scattering events as $l_{n,m}^{i}$. Here, $\mathbf{q} = \mathbf{k}_{out} - \mathbf{k}_{in}$, with $\mathbf{k}_{out}$ and $\mathbf{k}_{in}$ are wavevectors denoting light traveling from and towards a scattering collision, respectively. The total dimensionless momentum transfer and photon traveling path length of photon $n$ inside medium $m$ can then be written as $Y_{n,m} = \sum_{i=1} \left(\boldsymbol{q}_{\boldsymbol{n},\boldsymbol{m}}^{\boldsymbol{i}}\right)^2 / (2k_m^2)$ and $L_{n,m} = \sum_i l_{n,m}^{i}$, respectively[6], where the summation is performed over the number of scattering events that occur per photon. With the $\mathbf{q}_{n,m}^{i}$ and $l_{n,m}^{i}$ tracked for each photon via a Monte Carlo simulation, prior work has shown that the resulting field correlation can be calculated as,

$$G_1(\tau) = \frac{1}{N_p} \sum_{n=1}^{N_p} \exp\left(-\frac{1}{3} \sum_{m=1}^{M} Y_{n,m} k_m^2 \langle \Delta r_m^2(\tau) \rangle\right) \exp\left(\sum_{m=1}^{M} -\mu_{a_m} L_{n,m}\right). \tag{1}$$

As detailed in Ref.6, $M$ is the number of different tissue types within the phantom and $N_p$ is the number of detected photons at the location of interest (i.e., where we assume our detection fiber is placed). $k_m$ and $\mu_{a_m}$ are the wavenumber and absorption coefficient in medium $m$, and

$$\langle \Delta r_m^2(\tau) \rangle = 6 D_v \tau + v_b^2 \tau^2, \tag{2}$$

where $D_v$ is the Brownian diffusion coefficient and $v_b$ is the blood flow speed. For a background medium without random flow ($v_b$), it reduces to a Brownian motion. From our simulated field correlation curves, we generate a normalized field correlation,

$$g_1(\tau) = G_1(\tau)/G_1(0). \tag{3}$$

We then add realistic noise $\varepsilon_m(\tau)$ distributed over $\mathcal{E}(\tau)$, following the method developed by Zhou *et al.* in Ref. 7, to the $m^{th}$ curve at each time lag using the photon number that matches our experiments. After adding noise, we calculate a final averaged curve:

$$\overline{g_1}(\tau) = \frac{1}{M}\sum_{m}^{M}\widehat{g_m}(\tau) = g_1 + \varepsilon_m(\tau) \tag{4}$$

Here, $M$ denotes the number of SPAD used in the simulation to measure the DCS signal and is selected from 1 to 1024 to demonstrate the accuracy improvement using massively parallel measurements. Eq. (4) represents our final means of simulating DCS autocorrelation curves via Monte Carlo simulation. We stick with $g_1(\tau)$ here instead of $g_1(\tau)$ as the autocorrelations calculated from the Monte Carlo simulation are non-negative values; hence the Siegert relation here is a bijective function, $g_1(\tau) = \sqrt{g_2(\tau) - 1}$ , so the detection accuracy estimated from $g_1(\tau)$ can be directly implied to the detection accuracy estimated using the intensity autocorrelation $g_2(\tau)$.

In the presented simulations, we used a source-detector separation of 14.0 mm to match our experiments with 5 mm-thick phantoms. We simulated a tissue-like phantom with $\mu_a = 0.04\ \mathrm{mm}^{-1}$, $\mu_s = 13\ \mathrm{mm}^{-1}$, anisotropy $\mathrm{g} = 0.92$, and an index of refraction $n = 1.33$. We then embedded a cubic heterogeneity with blood-like optical properties: $\mu_a = 0.3\ \mathrm{mm}^{-1}$, $\mu_s = 82\ \mathrm{mm}^{-1}$, anisotropy $\mathrm{g} = 0.98$, and an index of refraction $n = 1.37$. The inclusion was placed 5 mm beneath the surface of this tissue phantom. The background optical properties were chosen to match the polystyrene microbead solution used in our experiments, and the optical properties for the heterogeneity are chosen from typical values for blood[6]. The Brownian motion coefficient $D_v$ of background medium is $44 \times 10^{-6}\mathrm{mm}^2/\mathrm{s}$ , fitted from experimental data. The Brownian diffusion coefficient $D_v$ and flow speed of the blood tissue $v_b$ is set to be $80 \times 10^{-6}\ \mathrm{mm}^2/\mathrm{s}$ and $3\ \mathrm{mm/s}$, respectively. These decorrelating coefficients also fall in the range of human blood tissue and blood flow[6,8].

Supplementary Figure 2(a) illustrates the tissue phantom used for the simulation, also mirroring the tissue phantom used in our phantom experiments, apart from the inclusion. Supplementary Figure 2(b) includes simulated autocorrelation curves for the homogeneous medium described above, with included noise. Here, we compare the correlation from a heterogeneous phantom perturbed by blood flow, with the one from homogeneous phantom with the same optical properties. The first row plots the curves averaged from 1, 9, and 1024 parallelized measurements. The second row plots the average of all curves from 1000 experiments. The blue curves are generated with the homogeneous phantom, while the red curves are generated with the heterogeneous phantom that includes a $3 \times 3 \times 3\ \mathrm{mm}^3$BF inclusion. It is clear that a similar trend of curve separability manifests itself as a function SPAD integration number $M$ as compared to our experimental data.

To investigate this in more detail, we attempted to classify the noisy BF simulation curves into one of two categories (heterogeneous versus homogeneous). We averaged 1000 autocorrelation curves for each homogeneous phantom with blood inclusion, and fitted the curved with $\overline{g_1}(\tau) = \hat{\beta}\exp(-\hat{\gamma}\tau)$. We report a perturbation is detected if $\hat{\gamma}_{heter} > \hat{\gamma}_{homo}$, where $\hat{\gamma}_{heter}$ and $\hat{\gamma}_{homo}$ are fitted $\hat{\gamma}$ for heterogeneous (with blood inclusion) and homogeneous phantoms, respectively, and report the detection accuracy for inclusions of different sizes with different number of averaged measurements.

The results of this experiment are shown in Supplementary Figure 2 (c). Here, we also plot our experimental accuracy of detecting DMD-induced perturbations of different sizes, along with the accuracy of detecting blood flow inclusions of different sizes. Here, the width of the DMD-induced perturbation area is 1.4, 2.1, 2.7, 3.4, and 4.1 mm, while the width of the simulated cubic blood flow region is 1.0, 1.5,

2.0, 2.5, and 3.0 mm. We note that the two curves exhibit the same trend of accuracy increase as a function of experimental area and simulated volume increase.

Supplementary Figure 2(d) similarly plots the detection accuracy from phantom experiment (red) and simulation of blood flow (blue) as a function of the number of parallelized measurements recorded (i.e., using a different number of SPAD array pixels), which also exhibit similar trends as observed experimentally. The side length of the experimental and simulated perturbations (area and volume) were selected to be 4.1 mm and 3.0 mm here, respectively. These curves roughly match those in Fig. 4(c) in the main text. These results suggest that our selected frequencies of DMD-induced perturbation (~kHz), which we picked intentionally, are well-suited for approximately matching the response expected by blood flow. These frequency values also fall into the optical frequency shift range introduced by blood flow as reported by various laser-doppler flowmetry studies[9,10]. As a final test, we also plot the simulated intensity autocorrelation curves for a BF phantom with perturbation side length of 4.1 mm for different blood flow speeds in Supplementary Figure 2(e). These curves show that decorrelation increases as a function of blood flow speed, as expected, and share a similar trend with the plots shown in Fig. 7(b) in the main text for our DMD-induced perturbation at different frequencies.

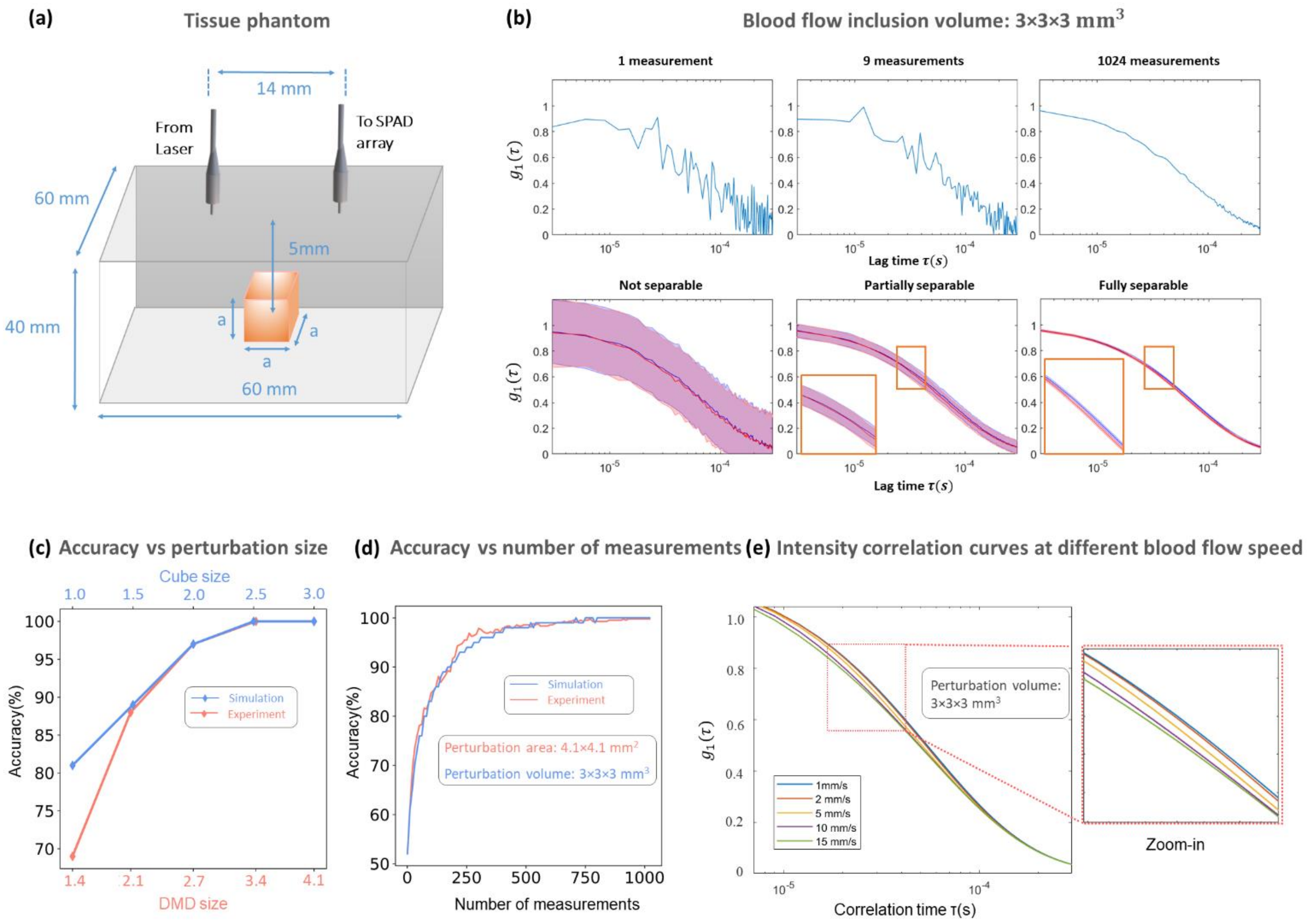


**Supplementary Figure 2.** **(a)** Illustration of tissue phantom structure investigated in blood flow simulation. **(b)** Simulated noisy autocorrelation curves. The first row plots the autocorrelation curves for the homogeneous medium described in Supplementary Note 2 averaged from 1, 9, and 1024 measurements. The second row plots all curves from 1000 experiments. Blue curves are generated with the homogeneous phantom, while red curves are generated with the heterogeneous phantom with the $3 \times 3 \times 3\ \mathrm{mm}^3$ blood like perturbation (all other properties unchanged). **(c)** Simulated and experimental detection accuracy for different sized perturbations. Blue line is simulated detection accuracy for cubic blood flow volumes of different side lengths. Red line is experimental detection accuracy for DMD-induced temporal perturbations of different side lengths. **(d)** Simulated and experimental detection accuracy for a fixed perturbation size (blood flow inclusion and DMD perturbation), produced by averaging over a different number of measurements $M$ from different SPAD pixels. Blue and red curves are simulated and experimental detection accuracies, respectively; the perturbation size for the simulation is $3 \times 3 \times 3\ \mathrm{mm}^3$ (BF inclusion volume), and is $4.1 \times 4.1\ \mathrm{mm}^2$ (DMD area) for the experiments. **(e)** Simulated intensity autocorrelation curves for a fixed perturbation size (same as used in (b)) with blood flow speed of 1,2,5,10, and 15 mm/s.

### Supplementary Figure 3:

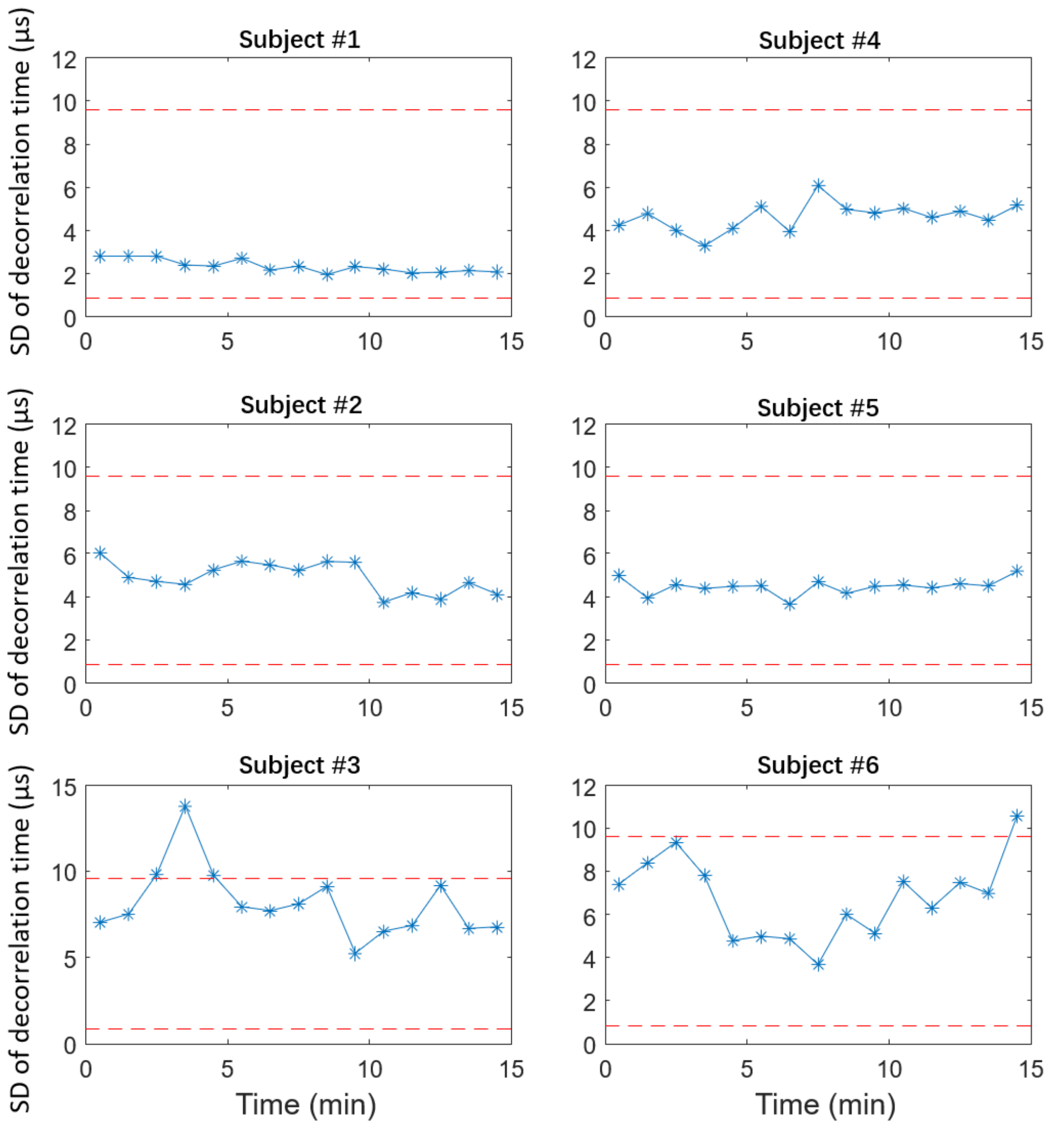


**Supplementary Figure 3.** The plot of the standard deviation (SD) of the measured decorrelation time, $\sigma_d$, during each measurement window for all 6 enrolled subjects in the human prefrontal cortex activation experiment. Red dash lines correspond to the 95% confidence intervals of $\sigma_d$.
